# An autologous cell-based therapeutic vaccine expressing IL6/1 fusokine drives robust anti-tumor response against ovarian cancer

**DOI:** 10.64898/2026.05.05.721149

**Authors:** Sejal Sharma, Rahul Das, Andrea Pennati, Catigan Hedican, Lisa Barroilhet, Manish S. Patankar, Jacques Galipeau

## Abstract

**Background:** Cytokines are immunomodulatory proteins that play central roles in regulating immune responses and represent attractive targets for cancer therapy. However, as single agents, cytokines have shown limited clinical benefit due to systemic toxicities and a short in vivo half-life. Our group has focused on engineering fusion cytokines (fusokines) that couple two cytokines into a single biologic to reprogram immune cell responses by enforcing non-canonical receptor engagement and signaling. A chimeric IL-6/IL-1β fusokine was engineered to test the hypothesis that enforced co-engagement of IL-6 and IL-1β signaling pathways would confer a gain-of-function phenotype in T cells and promote robust anti-tumor immunity. Here, we describe the immunomodulatory properties of IL6/1 fusokine and a method to deliver this fusokine to produce inhibition of ovarian tumor growth in a pre-clinical mouse model.

**Methods:** Lentiviral vectors encoding murine or human IL6/1 were designed using Vector Builder and expressed in either HEK293, CHO or ID8-F3 (p53^−/-^) cells depending on the downstream experiment to be conducted. IL6/1 expression was validated by ELISA and flow cytometry. Effects of human IL6/1 (hIL6/1) on T cell function (proliferation, memory phenotype, activation induced apoptosis) were monitored by flow cytometry. For in vivo studies, ID8-F3 murine ovarian cancer cells expressing mouse IL6/1 (mIL6/1) were administered intraperitoneally (I.P.) as a cell-based therapy to C57BL/6 female mice bearing established ID8-F3 luciferase tumors. Tumor progression was monitored by bioluminescence (BLI) imaging, and overall survival was evaluated.

**Results:** hIL6/1 significantly enhanced T cell survival and selectively promoted activation and expansion of CD45RO⁺ memory T cells. mIL6/1 expressing ID8-F3 cells (ID8IL6/1) demonstrated stable transduction and sustained cytokine secretion. In vivo, ID8IL6/1 cell therapy significantly reduced tumor growth and improved overall survival compared to control groups, with 2 of 8 mice achieving complete tumor clearance.

**Conclusion:** These findings indicate that IL6/1 fusokine enhances T cell survival and proliferation while promoting memory responses. Engineered cancer cells (ID8-F3) expressing mIL6/1 fusokine induced a strong anti-tumor response when delivered as a therapeutic vaccine in ovarian cancer mouse model.

**What is already known on this topic:**

- Fusokines are a class of bifunctional proteins designed to achieve synergistic immune modulation. Previous studies in our lab have shown fusokine exhibit gain-of-function immunomodulating activity. Individually, IL-6 and IL-1β are recognized for their roles in promoting T-cell proliferation and effector function. However, the potential for a fused IL-6/1 fusokine to reprogram the immune system and elicit a superior anti-tumor response *in vivo* in ovarian cancer model is not yet studied.

**What this study adds:**

- This study develops a novel fusion cytokine (fusokine), combining IL-6 and IL-1β, and demonstrate robust activation of T cells. In a preclinical ovarian cancer model, engineered cancer cells expressing IL6/1 used as a therapeutic vaccine showed significant tumor reduction and improved overall survival.

**How this study might affect research, practice or policy:**

- This study demonstrates that in comparison to individual cytokines, fusokines have greater potential to activate T cell function and when delivered as a cell therapy, achieve clear therapeutic efficacy in an ovarian cancer model. Further translational and clinical studies may enable the development of novel and more effective fusokine cell therapy approaches for patients with ovarian cancer.

## Introduction

Over the past decade, immunotherapy has emerged as a transformative modality in oncology, reshaping therapeutic paradigms across a range of malignancies [1, 2]. Current strategies encompass a diverse range of approaches, including antibody drug conjugates (ADCs) [3], bispecific antibodies, cancer vaccines, adoptive cell therapies, and immune checkpoint blockade, which have shown durable tumor control [1, 2]. Cytokines have also played a pivotal role as immunomodulatory agents; however, the clinical efficacy of these cytokines has been constrained due to short serum half-lives and dose-limiting toxicities [4, 5]. To overcome these limitations, our group has developed chimeric cytokines called fusokines that are chimeras of two functionally unrelated, yet potentially synergistic cytokines [6–8]. These fusokines bypass natural physiological regulation and can pharmacologically induce clustering of otherwise unrelated, yet activated, cytokine receptors [6]. This can result in unique and supraphysiological signals that ultimately lead to novel biological effects [9–11]. The first family of fusokines designed in our lab was a combination of GM-CSF as a common moiety with several interleukins like IL-2, IL-15,IL-21, IL-4, IL-7, and IL-9 to form fusion proteins GIFT2 [10, 12], GIFT21 [11], GIFT15 [13], GIFT4 [9], GIFT7 [8], and GIFT9 [14], respectively. Previous studies by our group have shown therapeutic benefits of fusokine in proinflammatory and anti-inflammatory conditions. For example, mice vaccinated with irradiated B16 melanoma cells secreting GIFT2 were protected against a subsequent melanoma tumor challenge [12]. In a therapeutic setting, more mice with established tumors were able to progress towards tumor-free state when treated with irradiated B16-GIFT2 cells, compared to those receiving cells secreting GM-CSF or IL-2 alone. GIFT2-treated mice also showed significantly increased NK cell numbers relative to controls. GIFT21 induces IL-21Rα-dependent STAT3 hyperactivation while concurrently inhibiting GMCSF mediated STAT5 signaling [11]. In vivo, GIFT21 secreting B16 melanoma cells are immune-rejected in syngeneic mice and confer a significant survival advantage in NOD-SCID mice. Furthermore, GIFT21 effectively eliminates IL-21Rα⁺ EL-4 lymphoma cells and prolongs survival in tumor-bearing mice. On the other hand, GIFT15 functions as an immunosuppressive agent in both B16 and U68 human glioblastoma models [13]. In a murine model of multiple sclerosis, intravenous infusion of GIFT15 treated B cells (GIFT15-Bregs) showed complete remission of disease with suppressed neuroinflammation and remyelination [15]. More recent study showed that delivery of GIFT7 tumor vaccine reversed age-related thymic atrophy and enhanced antitumor immunity in aged mouse models of glioblastoma by modulating T cell response [16]. Preclinical studies with this GM-CSF family of fusokines have shown a plethora of effects on T cells [8], B cells [9], NK cells [12], making them robust immunotherapeutic agents. Building on the success of GM-CSF based fusokines, our group has now developed a new class of non-GM-CSF fusokines incorporating additional cytokines to broaden their immunomodulatory potential. Interleukin 6 (IL-6) is a pleiotropic cytokine that regulates immune response by inducing activation, amplification, survival, and polarization of T cells [17]. Interlukin-1β (IL-1β) is mainly a pro-inflammatory cytokine that plays a critical role in the immune response by augmenting the expansion of antigen-primed CD4 and CD8 T cells in vivo, and activation of T cells by antigen-presenting dendritic cells [18, 19]. IL-6 signals through the JAK/STAT signaling pathways [20], whereas IL-1β primarily acts through the MAP kinase signaling pathways [21]. Given the central roles of IL-6 and IL-1β in T-cell biology, we engineered a novel fusion cytokine, IL-6/1, by physically linking the IL-6 and IL-1β proteins. Here, we report on the design and construction of this fusokine and characterize its immunomodulatory effects on immune cells, with a particular focus on T cells. We further show that delivery of the IL6/1 fusokine via engineered cancer cells significantly inhibits tumor growth and improves overall survival in a preclinical mouse model of ovarian cancer. The current manuscript is the first to study this fusokine as well as a novel platform of IL6/1 delivery and demonstrate its therapeutic efficacy in a cancer model. Additionally, ovarian tumors are generally considered immunologically “cold,” exhibiting limited T cell infiltration and poor responsiveness to immunotherapy [22, 23]. Therefore, we hypothesized that delivery of the fusokine as a cell-based therapy could modulate the tumor microenvironment (TME) to enhance immune activation and convert these tumors into an immunologically “hot” state. Data presented in this study provides a foundation for the use of IL-6/1, demonstrating its ability to activate immune responses and highlighting its potential as a novel therapy for ovarian cancer treatment.

## Methods

### Lentiviral vector design

IL-6 and IL-1β genes (human or mouse) were cloned downstream of the EF1α promoter in the pLVX lentiviral vector backbone (Vector Builder) to generate constructs encoding human or mouse IL6/1 variants. A puromycin resistance gene, driven by the mPGK promoter, was included for antibiotic selection. Lentiviruses expressing either the control vector (GFP or RFP), IL-6, IL-1β or IL6/1 constructs were produced using a standard calcium phosphate (Ca-P) transfection protocol in HEK293 cells. The transfection mixture contained the transfer plasmid, psPAX2 packaging plasmid, and pVSV-G envelope plasmid in a 5:2:1 ratio (transgene:psPAX2:pVSV-G). The purified viral particles were used to transduce multiple cell types for fusokine/cytokine production.

### Fusokine/Cytokine production

HEK293 cells were transduced with lentivirus carrying GFP or RFP control or IL6/1, IL-6 or IL-1β constructs using polybrene. 24 h post transduction, puromycin was added at 1 µg/mL and cells were selected for 7 days with intermittent media change. Stably transduced cells were used for protein production. Cells were seeded in 150 cm^2^ dishes at ∼30% confluence and cultured for 48-72 h. The conditioned media were collected and concentrated through a 30 kDa cutoff centrifugal filter. Concentrated media were aliquoted and stored at -80 °C.

### Western blot and ELISA

Cells were washed once with ice-cold PBS and lysed on ice using lysis buffer (Cell Signaling Technology) supplemented with 1mM PMSF (ThermoFisher). Lysates were sonicated on ice, centrifuged at 12,000xg for 15 min at 4°C and the supernatants were collected. Pierce™ 660nm Protein Assay Reagent (ThermoFisher) was used to determine protein concentration. Western blots were performed according to standard protocol. Primary antibodies were prepared in TBST +5% BSA according to the manufacturer’s recommendation. Images were taken using the Amersham Image Quant 800 imaging system (Cytiva). The primary antibodies anti-hIL-6 (CST, IL-6 (D3K2N) - 12153) and anti-hIL-1β (CST, IL-1β (D3U3E) - 12703) were used for the western blot.

mIL-6, mIL6/1, and mIL-1β secreted by ID8 cells were quantified with mouse IL-6 and mouse IL-1β ELISA kit (ThermoFisher).

### T cell isolation, activation, and proliferation

Human peripheral blood mononuclear cells (PBMCs) were isolated from healthy donor blood, and CD3⁺ T cells were purified by negative selection using the EasySep™ Human T Cell Isolation Kit (STEMCELL Technologies). For T cell activation, tissue culture-treated 12-well plates were coated with anti-CD3 antibody (BioLegend) at a concentration of 3 µg/mL in PBS and incubated overnight at 4 °C. The plates were washed three times with PBS prior to cell seeding. Purified T cells were plated at a density of 1 × 10⁶ cells/mL in RPMI-1640 medium supplemented with 10% fetal bovine serum (FBS), 1% penicillin-streptomycin-glutamine, 1 mM sodium pyruvate, non-essential amino acids (NEAA), and 10 mM HEPES. Soluble anti-CD28 antibody (BioLegend) was added to the culture medium at a final concentration of 5 µg/mL to provide co-stimulation. For proliferation assays, activated T cells were labeled with CellTrace™ Far Red proliferation dye (Invitrogen) according to the manufacturer’s protocol. Fluorophore minus-one (FMOs) for all flow cytometry data is in supplementary figure 1.

### Flow cytometry

Flow cytometric analyses were performed using an Attune NxT Flow Cytometer (Thermo Fisher Scientific). Antibody staining and washing were conducted in 1X PBS containing 5% FBS. To determine T cell memory phenotypes, cells were stained with anti-CD4, anti-CD-8, anti-CD45RA BV510 and anti-CD45RO PE antibodies (BioLegend) for 30 min at 4 °C, followed by one wash with FACS buffer (1x PBS containing 2% FBS and 0.09% sodium azide). For apoptosis assays, cells were washed once with Annexin V binding buffer (BioLegend) and incubated with Annexin V-APC/Fire™ 750 (BioLegend) for 20 min at room temperature in the dark. After incubation, cells were washed once and resuspended in Annexin V binding buffer containing DAPI (1:10,000) for dead cell exclusion. Samples were acquired on the flow cytometer, and data were analyzed using FCS Express 6 software (De Novo Software). In addition, ID8-F3 ovarian cancer cells transduced with the mIL6/1 fusion protein construct co-expressing Turbo Red fluorescent protein (TurboRFP) were evaluated by flow cytometry to confirm transduction efficiency.

### Generation of cytokines/fusokine expressing ID8-F3 ovarian cancer cells

Murine ovarian cancer ID8-F3 cells were seeded at 50-60% confluency in 150 cm² tissue culture plates and allowed to adhere overnight. The following day, cells were transduced with a lentiviral vector encoding either mouse IL-6, IL-1β, fusion IL6/1 or empty vector (EV), supplemented with 10 µg/mL polybrene to enhance transduction efficiency. Transduction was carried out by incubating cells with the viral supernatant at 37 °C in a humidified atmosphere containing 5% CO₂ overnight. After 24 h, the viral medium was replaced with fresh complete culture medium, and cells were maintained for an additional 48 h to allow for expression of the transgene. Transduced cells were then subjected to puromycin selection (2 µg/mL) for 24 h to enrich for stably transduced populations. Single-cell cloning was performed by limiting dilution to isolate monoclonal populations, which were subsequently expanded for downstream applications. Transduction efficiency was initially assessed by fluorescence microscopy for RFP expression encoded by the lentiviral construct and further quantified by flow cytometry. To confirm the functional expression of the mIL6/1 fusion protein, ELISA was performed on conditioned media collected from transduced ID8 cell cultures to quantify secreted mIL-6 and mIL-1β levels, as a proxy for IL6/1 expression.

### Animal housing and treatment experiments

All animals used in experiments were maintained in accordance with institutional IACUC guidelines established at the University of Wisconsin-Madison (Protocol number: M006730). Healthy 6-8 weeks old immunocompetent female C57BL/6 mice used for research were obtained from Research Animal Resources and Compliance (RARC). To mitigate potential confounders, all experimental cohorts were maintained under identical conditions and timelines to minimize environmental effects. All mice were contained in a temperature-controlled animal room maintaining a 12-h light and dark cycle with ad libitum access to food and water.

For all *in vivo* experiments, cultured cells were harvested by trypsinization and resuspended to obtain a single-cell suspension. Cell viability was assessed using trypan blue exclusion, and live cells were counted using a hemocytometer. The desired number of viable cells were then diluted in 100 µL sterile phosphate-buffered saline (PBS). Under aseptic conditions in a biosafety cabinet, mice were anesthetized, and the cell suspension was administered intraperitoneally (IP) using a 26-gauge needle and a syringe. ID8-F3 cells stably expressing luciferase (ID8 Luc^+^) were used to monitor tumor progression and quantify tumor burden via in vivo bioluminescence imaging system, the Lago Spectral imaging system (Spectral Instruments Imaging). Imaging data were analyzed using Aura imaging software (Spectral Instruments Imaging).

### Statistics

Data are presented as mean ± SEM. Statistical analyses were performed using one-way ANOVA followed by Dunnet’s post-hoc tests or Student’s *t*-tests, as appropriate. A p-value of less than 0.05 was considered statistically significant. For flow cytometry experiments, each graph represents data from an individual PBMC donor, as indicated in the figures. A total of three independent PBMC donors were used for each flow cytometry experiment (*n* = 3; biological replicates).

For in vivo experiment, tumor progression was quantified by measuring bioluminescence (BLI) using the Lago (Spectral Instruments) system. We established baseline tumor growth and then assigned mice to experimental groups without blinding. Tumor progression was monitored via longitudinal BLI; growth was quantified by normalizing luminescence signals to the baseline values measured for each mouse prior to start of treatment. Statistical comparisons between the treatment groups at individual time points were conducted using a multiple comparison test using GraphPad Prism 10. Animals reaching predefined endpoints, including signs of pain and distress like hunched posture, tumors impeding mobility, respiratory distress were humanely euthanized in compliance with approved animal care protocol. The mice were monitored for upto 120 days after treatment initiation to plot survival. The overall survival curve was plotted using Kaplan-Meyer plot and median survival times were calculated. Statistical significance was defined as p < 0.05.

## Results

### Generation and biochemical characterization of the chimeric human IL6/1 (hIL6/1) fusion cytokine

To combine the hIL-6 and hIL-1β cytokines in a single reading frame, we used the full-length coding sequence of each cytokine, including the N-terminal secretion signal, from the human IL-6 gene. To enable constitutive secretion of IL-1β, the N-terminal propeptide sequence (amino acids 1–116) was removed. This modification bypasses the native secretion constraints of wild-type IL-1β, as the propeptide region inhibits secretion. As shown in Fig. 1A, two fusion constructs were generated: an IL-6–IL-1β configuration (N- to C-terminal orientation) in which the cytokines were linked by a 3×EAAAK linker, and an IL-1β-IL-6 configuration in which a synthetic α-lactalbumin secretion signal peptide (MMSFVSLLLVGILFHATQA) was added at the N terminus to facilitate secretion. Both constructs were subcloned downstream of a full-length EF1α promoter in a second-generation lentiviral vector encoding a NeonGreen fluorescent reporter and puromycin resistance cassette. The final hIL-6/1 fusokine cDNAs encoded single polypeptide chains with a predicted molecular weight (non-glycosylated) of approximately 42 kDa, which was confirmed by Western blot analysis (Fig. 1B). Human HEK293 cells were transduced with lentiviral vectors carrying the fusokine constructs and generated clonal cell lines expressing hIL6/1. Cells were expanded and media collected at 24 h and 48 h intervals. Culture supernatants were subjected to SDS-PAGE/Western blot using monoclonal antibodies against hIL-6 or hIL-1β. As shown in Fig. 1B, IL-1β antibody detected both forms of fusokine, with higher levels detected after 48 h culture. In contrast, cell lysates showed a much smaller amount of fusokine accumulation, indicating that most of the protein is secreted in the media. On the other hand, IL-6 antibody detected the fusokine with IL-6-IL-1β orientation but not in IL-1β-IL-6 orientation; indicating that the epitope structure of IL-6 is retained only in IL-6-IL-1β orientation. Therefore, we decided to use IL-6-1β (IL6/1) oriented fusokine.

**Figure 1.**
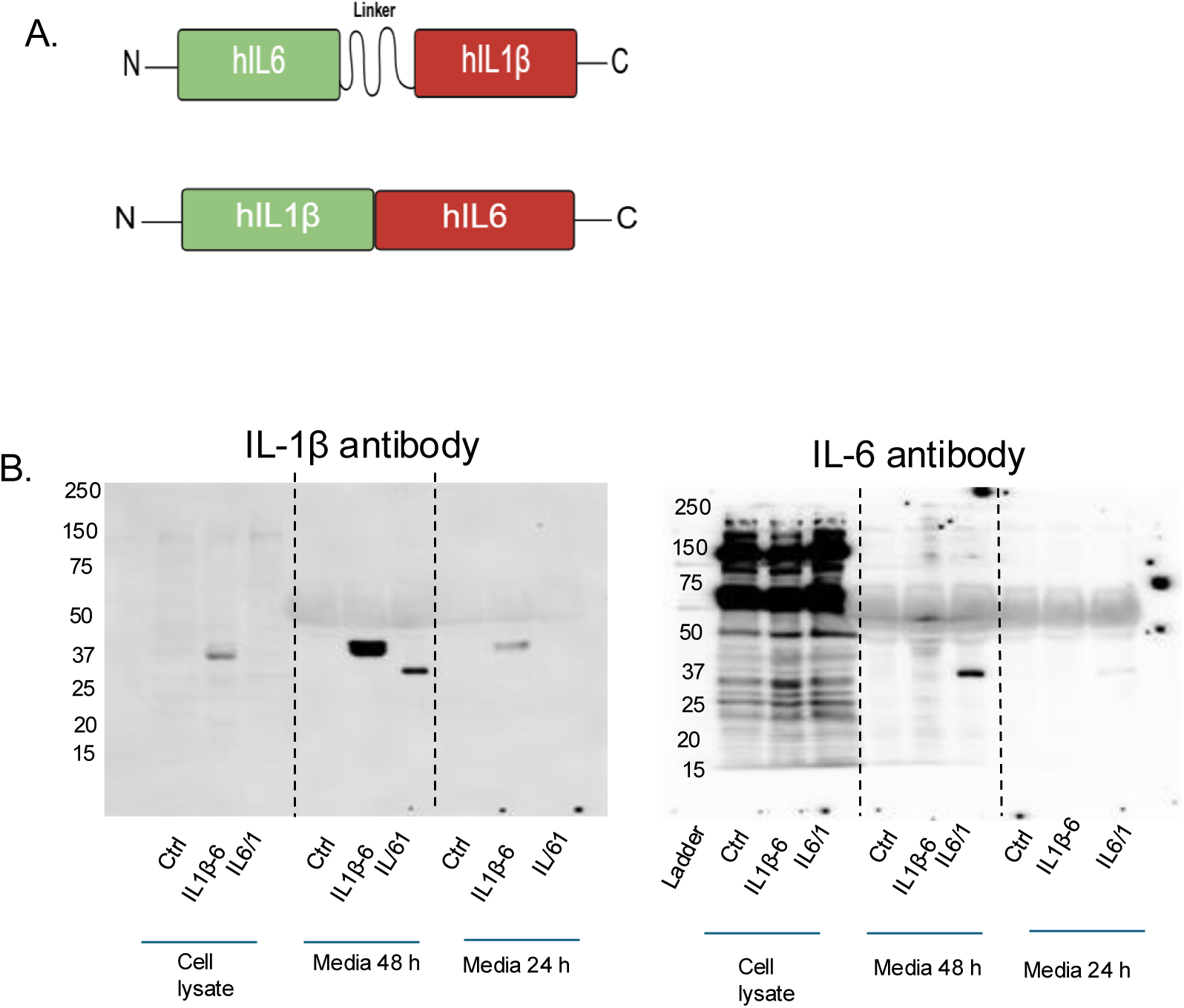
Characterization of chimeric hIL6/1 fusion cytokine: (A) Spatial orientation of components and amino acid sequences of IL6/1 and IL-1β-IL-6 fusion cytokines are shown. Green – IL-6, Red - IL-1β; the pro-IL-1β peptide was deleted from IL-1β sequences and lactalbumin secretion sequence was used in IL-1β -IL6 orientation. (B) Western blots of IL-6-IL-1β constructs with opposing orientations. HEK293 cells were stably transduced with constructs carrying hIL6/1 (IL-6-IL-1β orientation) or IL-1β-IL-6 (IL-1β -IL6 orientation). Conditioned media were collected from stable cell culture post 24 or 48 h of culturing. After 48 h, cells were lysed and media/ lysates were resolved by 10% reducing denaturing SDS-PAGE. Then protein was transferred to nitrocellulose membrane and probed with the specific antibodies mentioned.

### hIL6/1 fusokine potentiates proliferation of human T cells

Studies have shown that both IL-6 and IL-1β can act as proinflammatory cytokines to enhance T cell activation and proliferation [17, 24, 25]. We first tested if fusokine IL6/1 can promote stronger T cell proliferation compared to its individual counterparts. Human T cells isolated from healthy donor PBMCs were cultured in the presence of IL-2 and anti-CD3 anti-CD28 antibodies, which induce robust T cells proliferation by simultaneously engaging CD3 and CD28 receptors on the surface of the T cells. To track the rate of proliferation of T cells, CellTrace^TM^ Far Red (CTFR) proliferation due was used which binds to the intracellular proteins and the fluorescence intensity is approximately halved every successive cell division.

Next, we treated these cells for 4 days in the presence of hIL-6, hIL-1β, hIL-6 + hIL-1β, hIL6/1 and conditioned media control. Treatment with hIL6/1 increased the percentage of proliferating CD4⁺ (hIL6/1 40.4% vs hIL-6 26.6% vs hIL-1β 26%) and CD8⁺ (hIL6/1 47.1% vs hIL-6 31.8% vs hIL-1β 32.7%) T cell compared to controls (Fig. 2A-E) and a decrease in non-proliferating fraction of CD4⁺ (hIL6/1 15.92% vs hIL-6 37.5% vs hIL-1β 38.4%) and CD8⁺ (hIL6/1 9.37% vs hIL-6 23.2% vs hIL-1β 23.8%) T cell compared to controls (Fig. 2A-E). Overall, this indicates that the hIL6/1 treatment promotes the proliferation of a subset of T cells that may be refractory to rapid proliferation. To test whether these T cells show an enhanced memory-like phenotype, we stained these cells with anti-CD45RO and anti-CD45RA antibodies. As expected, hIL6/1 treated T cells showed increased percentage of CD45RO in the fraction of CD4⁺ (hIL6/1 5.2% vs hIL-6 3.1% vs hIL-1β 2.9%) T cells (Fig. 3B, D) and CD8⁺ (hIL6/1 1.8% vs hIL-6 1.1% vs hIL-1β 1.2%) (Fig. 3C, E) T cells, indicating that hIL6/1 indeed induces the proliferation of T cells with a memory-like phenotype. In agreement with this hypothesis, CD45RA staining did not show any appreciable difference between the different treatment groups (Fig. 3A-C).

**Figure 2.**
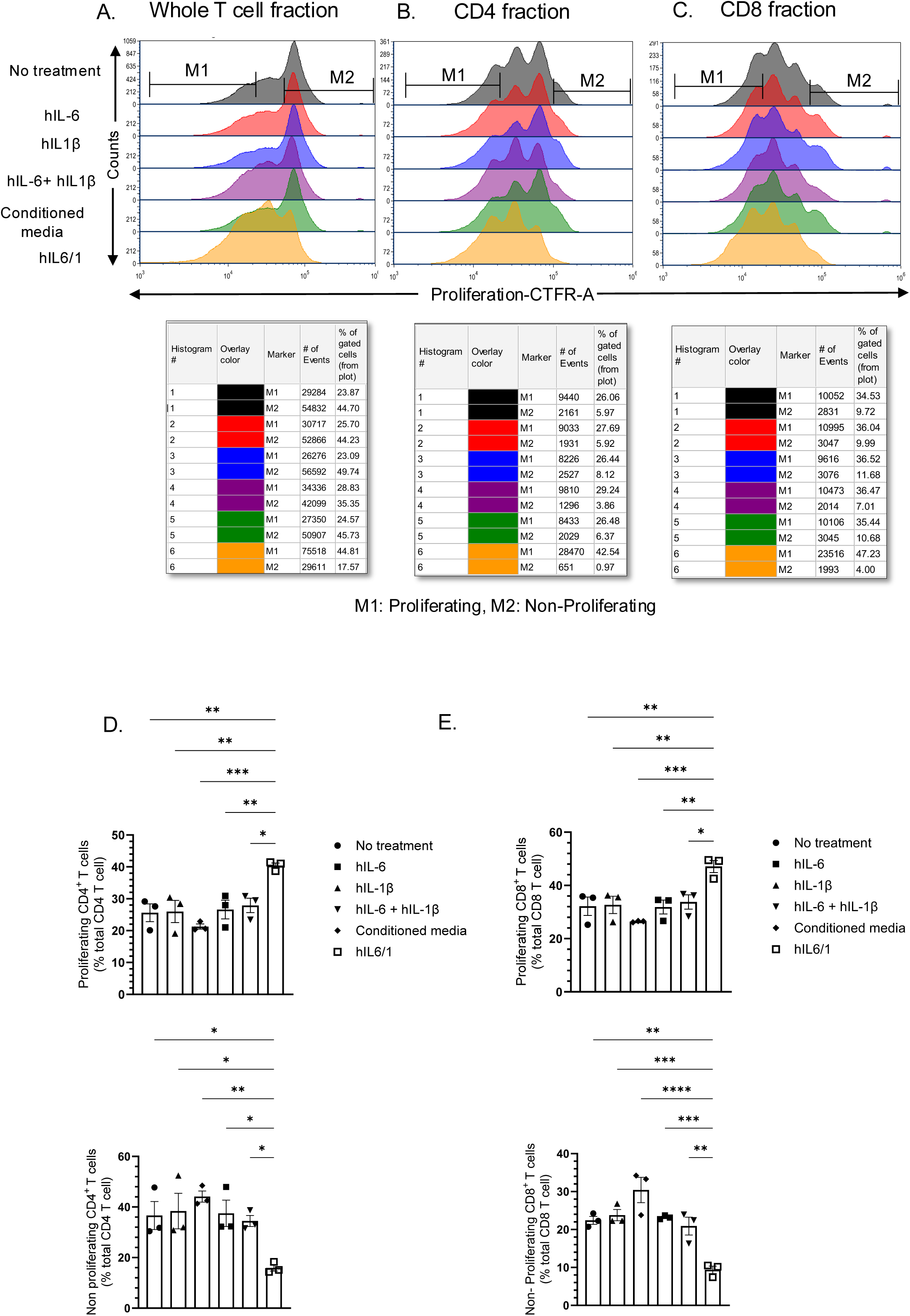
hIL6/1 induces selective proliferation of human PBMC derived T cells: Human peripheral blood mononuclear cells (PBMC) from healthy donors were isolated and CD3^+^ T cells were purified. Cells were labeled with CellTrace™ Far Red (CTFR) (CTFR) dye to track proliferation. Then cells were treated with anti-CD3 anti-CD28 monoclonal antibodies in combination with IL-2 cytokine for ∼96 h to induce proliferation. Cells were treated with hIL-6, hIL-1β, a combination of hIL-6 and hIL-1β or hIL6/1 fusokine (produced in HEK293 cells). Then cells were stained with anti- CD4 and CD8 antibodies and analyzed using flow cytometry. (A) Whole T cell population, (B) CD4 fraction, and (C) CD8 fraction. Quantitative analysis for measurement of proliferating and non-proliferating (D) CD4 and (E) CD8 T cells using One-way ANOVA followed by Dunnet’s multiple comparison is shown (N=3)

**Figure 3.**
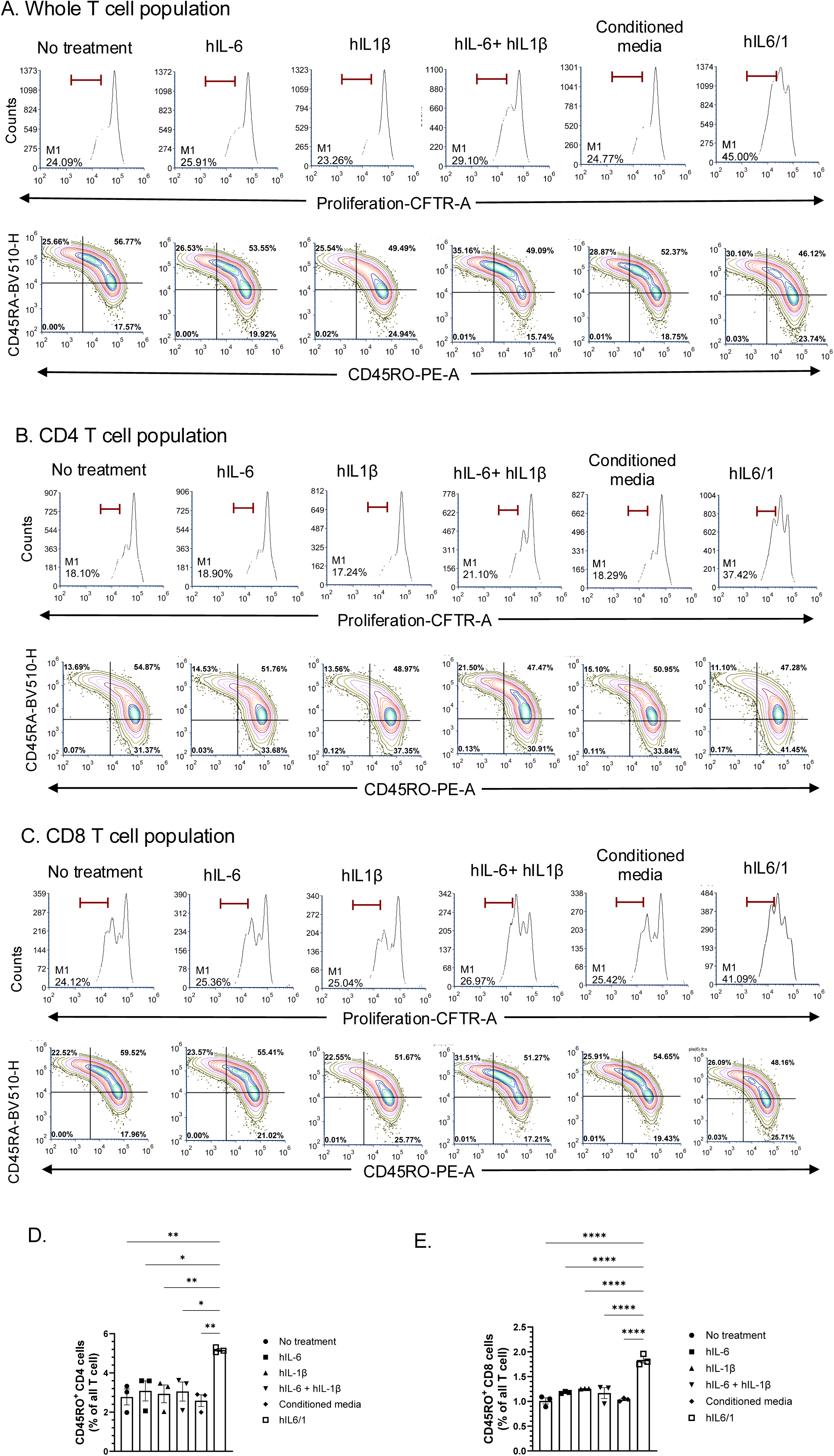
hIL6/1 treatment promotes proliferation of CD45RO memory T cell proliferation: Human peripheral blood mononuclear cells (PBMC) from healthy donors were collected and CD3^+^ T cells were purified. Cells were labeled with CTFR dye to track proliferation. Then cells were treated with anti-CD3 anti-CD28 monoclonal antibodies in combination with IL-2 cytokine for ∼96 h to induce proliferation. Cells were simultaneously treated in the presence of IL-6, IL-1β, a combination of IL-6 and IL-1β or IL6/1 fusokine (produced in HEK293 cells). Then cells were stained with anti- CD4, CD8, CD45RA, and CD45RO antibodies and analyzed using flow cytometry. (A) Whole T cell population, (B) CD4 fraction, and (C) CD8 fraction. (D, E) A representative analysis performed using One-way ANOVA followed by Dunnet’s multiple comparison is shown (N=3).

### hIL6/1 prevents activation induced apoptosis in human T cells

T cells undergo activation induced apoptosis, where a significant portion of the T cells receiving a stimulatory signal from CD3 and a co-stimulatory signal from other receptors undergo apoptosis. hIL-6 and hIL-1β are known to promote T cell survival under various conditions [26–29]. To test whether hIL6/1 has any protective role under activating conditions, we treated human PBMC derived T cells with anti-CD3/anti-CD28 beads, which increases activation and proliferation of T cells by crosslinking glycosylated surface proteins, including the T cell receptor (TCR). Next, we treated these anti-CD3/anti-CD28- activated cells with the conditioned media control, individual cytokines, their combination, or hIL6/1 fusokine. The anti-CD3/antiCD28 activated T cells upon hIL6/1 treatment showed significant increase in percentage of non-apoptotic T cells (hIL6/1 66.6% vs hIL-6 51.5% vs hIL-1β 50.3%) (Fig. 4B, E) and subpopulations of CD4^+^ and CD8^+^ T cells showed a significant increase in percentage of non-apoptotic (CD4: hIL6/1 72.2% vs hIL-6 58.0% vs hIL-1β 56.8%), (CD8: hIL6/1 66.6% vs hIL-6 55.4% vs hIL-1β 53.4%) (Fig. 4C, D-G). hIL-6, hIL-1β or their combination only marginally increased cell viability (Fig. 4E-G), but IL6/1 fusokine treatment significantly increased cell viability and prevented apoptosis, indicating that compared to the component cytokines, IL6/1 showed a gain of function phenotype whereby it prevents activation induced T cell apoptosis. Mechanistically, we observed that hIL6/1 treatment on T cells increased phosphorylation of STAT3 compared to other treatment conditions. In contrast, no difference was observed in other pSTAT proteins or total proteins, indicating that hIL6/1 selectively promotes STAT3 phosphorylation (Supplementary Fig. 2A). However further studies are required to fully understand the downstream consequences of this protein activation following hIL6/1 treatment.

**Figure 4.**
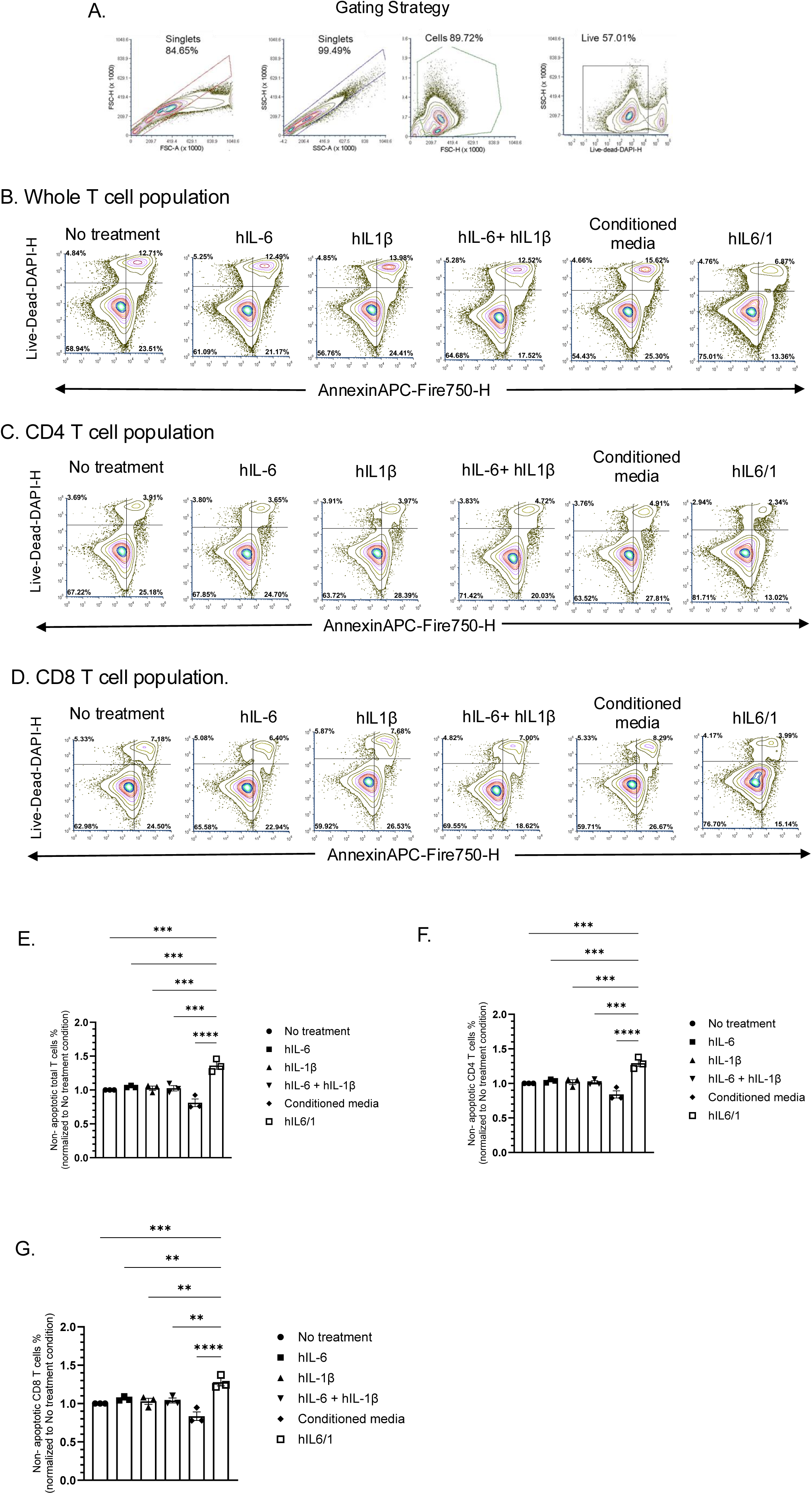
hIL6/1 protects T cells from activation induced apoptosis: Human PBMC derived CD3^+^ T cells were induced to proliferate by CD3/CD28 in the presence of equivalent amounts of hIL-6, hIL-1β, a combination of hIL-6 and hIL-1β or hIL6/1 fusokine (5 ng/ml). After 96 h of culturing, cells were stained with Annexin V and DAPI to determine the degree of apoptosis using flow cytometry: (A) Gating strategy, (B) Whole T cell population, (C) CD4 fraction, (D) CD8 fraction. A representative analysis for percentage of (E) non-apoptotic total T cells, (F) non-apoptotic CD4 T cells and (G) non-apoptotic CD8 T cells was performed using One-way ANOVA followed by Dunnet’s multiple comparison is shown (N=3). Specifically, the data was normalized to the no treatment group for reducing baseline variability.

### Engineering ID8-F3 cells to express individual cytokines or IL6/1 for in vivo experiments

ID8-F3 cells are murine ovarian cancer cells syngeneic to C57BL/6 mice commonly used in in vivo studies. To study the therapeutic effects of IL6/1 for ovarian cancer, we utilized an autologous cell therapy approach as a delivery system. We first designed a mouse IL6/1 (mIL6/1) fusokine construct. The mouse IL-6 (mIL-6) and mouse IL-1β (mIL-1β) cytokines were linked by an EK1GS2EK2 linker in the lentiviral vector design, as shown in Fig. 5A. The 3D structure of IL6/1 was predicted and constructed using AlphaFold Server structure prediction software (Fig. 5B). This construct was subcloned downstream of a full length EF1α promoter in a vector containing TurboRFP fluorescent marker and ampicillin antibiotic resistance gene. The vector was transfected in HEK293 cells to prepare viruses for transduction. The individual lentiviruses were transduced in ID8-F3 cells and transduction efficiency were qualitatively monitored by detecting TurboRFP expression by fluorescence microscopy (Fig. 5C). Quantitative measurements of transduction efficiency was performed using flow cytometry revealing ∼85% in ID8 empty vector (ID8EV) control cells, ∼70% in ID8IL6/1 transduced cells, ∼98% in ID8IL-6 cells and ∼95% in ID8IL-1β (Fig. 5D, E). Media collected from the transduced ID8-F3 cells were measured for cytokine secretion and showed a robust increase in mIL-6 (Fig. 5F) and mIL-1β (Fig. 5G) in ID8 cells expressing mIL6/1. ID8-F3 cells that were transduced with mIL-6 and mIL-1β showed only the expression of these respective cytokines (Fig. 5F, G). These results validated the expression of cytokines/IL6/1 in ID8-F3 ovarian cancer cells leading us to use these cell lines for the in vivo experiments described below.

**Figure 5.**
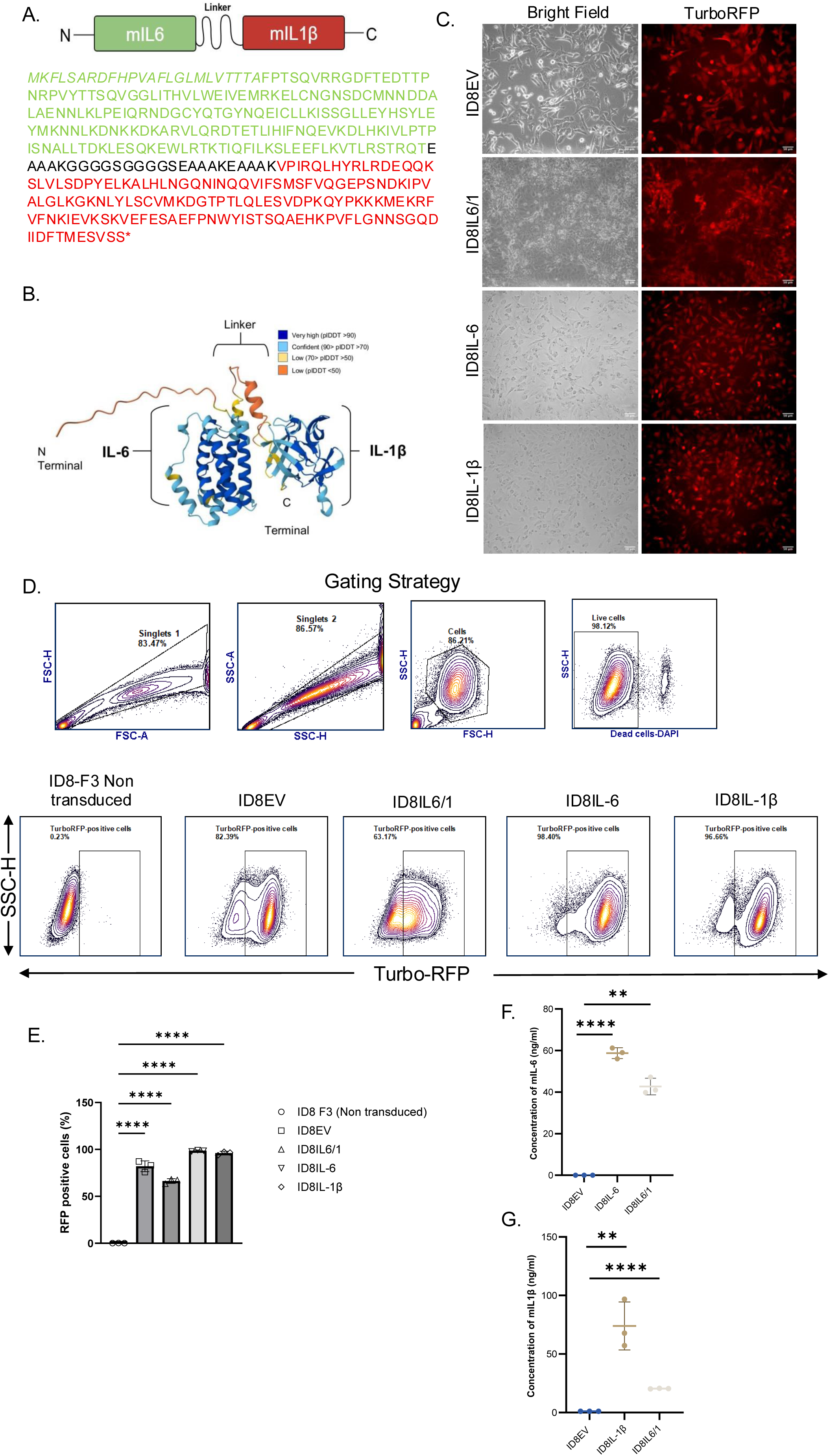
Engineered and validated ID8-F3 cells expressing cytokines and mIL6/1 fusokine: (A) Spatial orientation of components and amino acid sequence of mIL6/1 fusion cytokine is shown. Green – mIL-6, Red- mIL-1β; Note that pro-IL-1β peptide was deleted from IL-1β sequence. (B) Predicted ribbon structure of mouse IL6/1 fusokine: Secondary ribbon structure obtained using AlphaFold Server prediction software. (C) Lentiviral transduction efficiency of ID8 ovarian cancer cells post transduction. Scale bar = 50 um. (D) Gating strategy and phenotypic characterization of ID8 cells transduced with lentiviral vectors expressing mIL-6, mIL-1β, mIL6/1, EV (empty vector). (E) Data represents percentage of TurboRFP positive cells to determine successful transduction by flow cytometry. (F, G) Cell culture supernatants from ID8 cells expressing mIL-6, mIL- 1β or both mIL-6 and mIL-1β. Data are expressed as mean ± SEM from three independent experiments (N=3).

### Therapeutic administration of engineered cancer cells secreting mIL6/1 delays tumor growth and improves overall survival in a high-grade serous ovarian cancer model (HGSOC)

HGSOC is an immunological “cold” tumor with a complex TME [22, 23]. Based on our previous data, we hypothesize that delivery of IL6/1 will provide a strong anti-tumor response by modulating T cell activity. Also, studies conducted by Shireman.et.al showed that delivery of engineered cancer cells expressing GIFT7 fusokine reversed age-related thymic atrophy and enhanced antitumor immunity in aged mouse models of glioblastoma [16]. Using a similar IL6/1 delivery, we engineered and validated ID8-F3 cancer cells expressing IL6/1 or IL-6 or IL-1β (Fig. 6) and designed a therapeutic vaccine pipeline to test IL6/1 efficacy in an in vivo ovarian cancer model (Fig. 6A).

**Figure 6.**
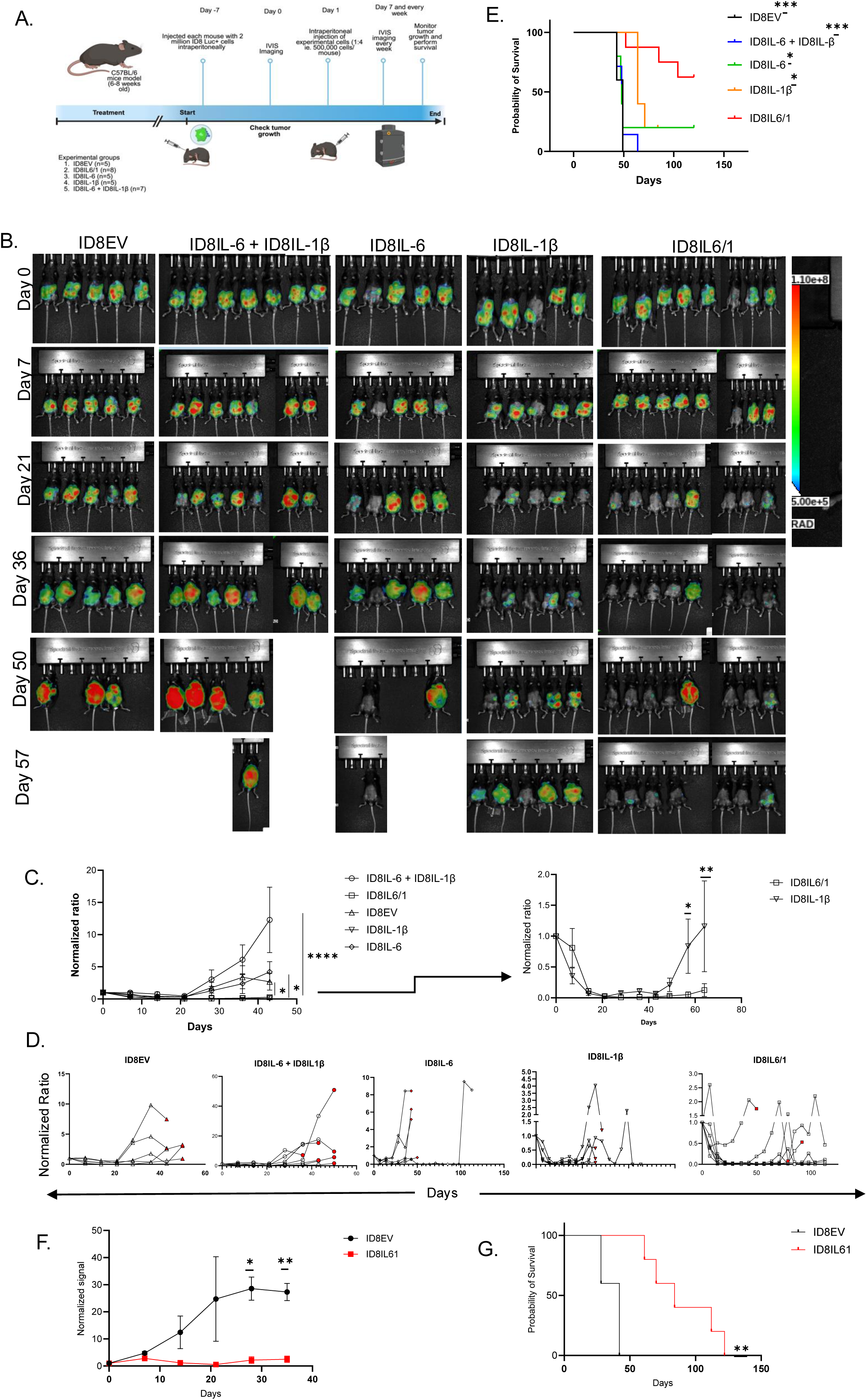
mIL6/1 engineered cell therapy shows therapeutic benefit and improves overall survival in ovarian cancer mouse model. (A) Schematic representation of the experimental design and treatment schedule for in vivo study of mIL6/1 engineered cell therapy in ovarian cancer mouse model. (B) Tumor growth was quantitatively analyzed using bioluminescence (BLI) signals obtained from LAGO imaging system at a week after tumor inoculation and subsequent weeks. Longitudinal images of a representative mouse from each treatment group were presented from a week after tumor inoculation before treatment and subsequent weeks. (C) Total emission signal (photons/sec) were calculated by Aura software and normalized to the signal before start of treatment. Statistical significances were analyzed using Two-Way ANOVA followed by Tukey’s multiple comparison tests. Data were indicated as the mean ± SEM or Student *T* tests. (D) Spaghetti plot for each mouse in all experimental groups are shown. (E) Kaplan-Meier survival curves for mIL6/1 cell therapy. Mice were considered dead if found dead in their cage or predefined endpoints due to health conditions or abdominal distension. (F) Tumor growth as shown by luciferase signal measured using Bioluminescence (BLI) method and the signal measured (photons/sec) were calculated by Aura software and normalized to the signal before start of treatment. Statistical significances were analyzed using Two-Way ANOVA followed by Tukey’s multiple comparison tests. Data are presented as the mean ± SEM. (G) Kaplan-Meier survival curves for ID8IL6/1 vs ID8EV cell therapy. Mice were considered dead if found dead in their cage or predefined endpoints due to health conditions or abdominal distension.

Once the ID8-Luc tumor was established in the peritoneal cavity, mice were treated with 500,000 cells of ID8IL6/1 or Empty Vector (ID8EV) or ID8IL-6 only or ID8IL-1β only or ID8IL-6 + ID8IL-1β and monitored tumor growth and survival. After a single dose of cell therapy, we observed a significant decrease in tumor burden with mice treated with ID8IL6/1 cell therapy with 2 out of 8 mice completely tumor free (Fig. 6B-D). We also observed significantly better overall survival in ID8IL6/1 cell therapy group compared to all the other controls (Fig. 6E). Specifically, the tumor growth kinetics in ID8IL-1β induced a reduction in tumor burden comparable to ID8IL6/1. However, the inhibitory effect of ID8IL-1β was transient, the ID8IL6/1 group showed a significantly more sustained suppression of tumor growth over the course of study suggesting a synergistic effect of IL6/1. Through repeated in vivo experiments comparing ID8IL6/1 (n=5) vs ID8EV (n=5), we consistently observed a decreased tumor burden in mice treated with ID8IL6/1 compared to controls, alongside a significant increase in overall survival (Fig. 6F, G, Supplementary Fig. 2B). These results suggest that the IL6/1 fusokine when delivered as a cell therapy in a therapeutic vaccine approach, reduces tumor growth and extends overall survival. Collectively, these findings demonstrate the therapeutic relevance and efficacy of the ID8IL6/1 therapeutic vaccine strategy in a clinically relevant context.

## Discussion

IL-6/1 is a novel class of fusokine designed in our lab, and for the first time, we have demonstrated its mechanisms of action and effects on T cells with an anti-tumorigenic role in the ovarian cancer model. In this study, we designed a synthetic fusion cytokine, namely IL6/1, by linking Interleukin 6 (IL-6) and Interleukin-1β (IL-1β), both of which are known to elicit potent T cell modulatory activities. For the first time, we show the proliferative and anti-apoptotic effects of IL6/1 on primary human T cells in vitro and it’s anti-tumorigenic role in a murine ovarian cancer model. We also show that the orientation of the fusion protein plays a role in retaining the epitope structure of hIL-6 and is not subjected to protease-mediated degradation.

Due to specific antigen-directed cytotoxic function, T cells have become a central focus for anti-cancer cell-therapy that led to the development of CAR-T cells, which have produced significant responses in patients with B-cell leukemia or lymphoma [30]. However, T cell (including CAR-T cell) mediated anti-tumor activity suffers from several limitations, including limited lifespan of effector cells, limited antigen specificity, and poor tumor infiltration [31]. In this study, we have shown that hIL6/1 displays novel properties that enhance T-cell activation and survival compared with IL-6, IL-1β, or their combination. Memory T cells arise from antigen-exposed, proliferatively activated effector T cells. These memory T cells, characterized by the surface expression of CD45RO, persist for a much longer period and can coordinate protective immune responses following re-exposure. It has been reported that T cells with memory phenotypes exhibit superior in vitro and in vivo anti-tumor function [32]. Our results indicate that hIL6/1 fusion cytokine promotes memory-like-phenotype T cells. This effect was not observed in IL-6, IL-1β or a combination of IL-6 and IL-1β treated T cells, indicating an acquired gain of function by the fusokine. Upon receptor mediated activation, T cells undergo expansion followed by a contraction phase, during which many activated T cells undergo apoptosis. Enhancement of T cell survival can be beneficial in CAR-T cell therapy as well as cancer immunotherapy. Here we show that although IL-6, IL-1β or their combination showed a modest increase in activated T cell survival, hIL6/1 fusokine further enhanced T cell survival upon activation, indicating that this fusion protein may be suitable for T cell proliferation in in vitro settings where CAR-T cells are expanded before re-introducing into the patient. Overall, our study shows that hIL6/1 fusokine is a novel and potent tool for *ex vivo* expansion of activated T cells and selectively promotes memory T cell activation, which may hold therapeutic potential in cell-based immunotherapy of cancer.

Ovarian cancer is characterized as a “cold” tumor with a profoundly immunosuppressive microenvironment [22, 23]. We hypothesized that expression of a fusion cytokine, mIL6/1, by tumor cells could overcome this suppression by enhancing T cell activation, as demonstrated in our in vitro studies. Indeed, ID8-F3 cells expressing the mIL6/1 fusokine significantly reduced tumor growth and improved overall survival. In contrast, ID8IL-1β showed only transient reduction in tumor growth. The sustained anti-tumor response observed in the ID8IL6/1 may be due to the gain-of-effect of IL-6 and IL-1β unique from its individual responses. Together, these cytokines prime a more robust and durable pro-inflammatory T cell response compared to ID8IL-1β alone. Furthermore, the pharmacokinetic profile of the ID8IL6/1 likely plays a role; given that its molecular weight is approximately double that of monomeric cytokines, which may contribute to an extended *in vivo* half-life, allowing for extended therapeutic signaling. This antitumor effect is likely mediated by both adaptive and innate immune responses, resulting in enhanced recognition and engagement of tumor cells by immune effectors. Future studies will be performed to study the exact mechanism of action of IL6/1 and its direct effects on immune cell population in vivo. Nevertheless, our studies provide an insight into developing new immunotherapeutic drugs that can activate immune cells and acts as a strong anti-cancer agent in various cancer types.

## Presented at

Part of this work was presented at SITC 2024, AACR 2024 and AACR 2025 (Advances in Ovarian Cancer Research).

## Acknowledgments

The authors would like to thank University of Wisconsin-Madison RARC staff and veterinary team for their assistance with animal housing.

## Author contribution

S.S contributed to writing of manuscript and concept for in vivo work, ran experiments, analyzed and interpreted the data. R.D contributed to the idea and concept of fusokine and ran experiments, analyzed and interpreted the in vitro data.

A.P. contributed to interpretation of data and edited the manuscript. C.H. contributed to help with in vivo assays. L.B contributed to the concept of the work, interpretation of the data and edited the manuscript. M.P. and J.G contributed to the concept of the work, interpret data and supported to writing the manuscript. All the authors reviewed the manuscript and approved the final version.

## Patient consent for publication

All patient samples and data were de-identified, and no identifiable information is included in this manuscript. Therefore, specific consent for publication was not required.

## Ethics Approval

Human samples were obtained with written informed consent under protocols approved by the Institutional Review Board of the University of Wisconsin–Madison (IRB Protocol No. 2019-0865). All animal experiments were conducted in accordance with Institutional Animal Care and Use Committee (IACUC) guidelines at the University of Wisconsin-Madison (IACUC Protocol No. M006730).

## Data availability statement

Data are available in a public, open access repository and also available upon request.

