## Supplementary Figures for "An autologous cell-based therapeutic vaccine expressing IL6/1 fusokine drives robust anti-tumor response against ovarian cancer"

Supplementary Figure 1:

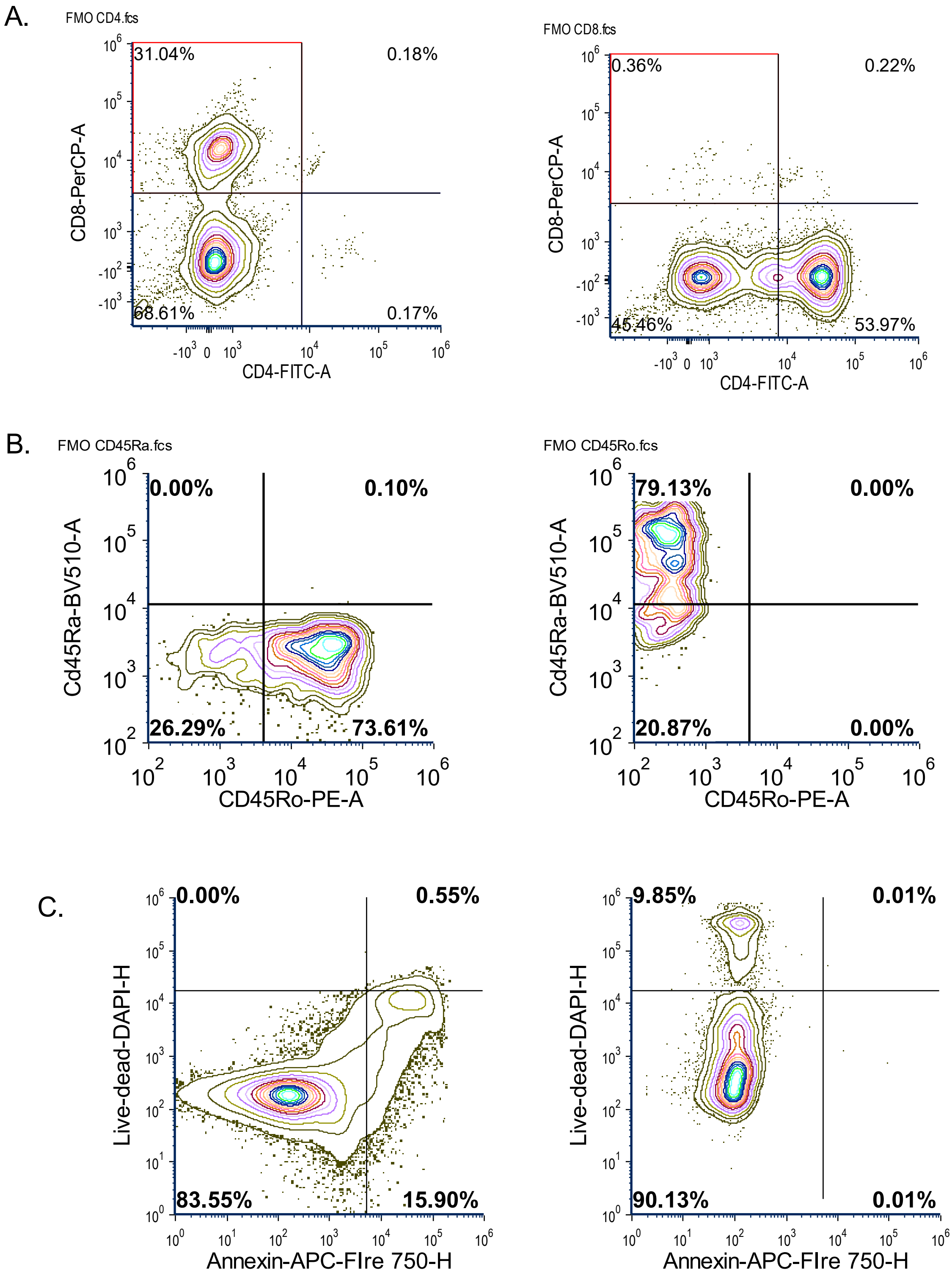


**Supplementary Figure 1:** (A) FMOs for CD4 and CD8 T cell proliferation data. (B) FMOs for CD45Ra and CD45Ro T cell data. (C) FMOs for DAPI and Annexin v for T cell apoptosis data.

Supplementary Figure 2:

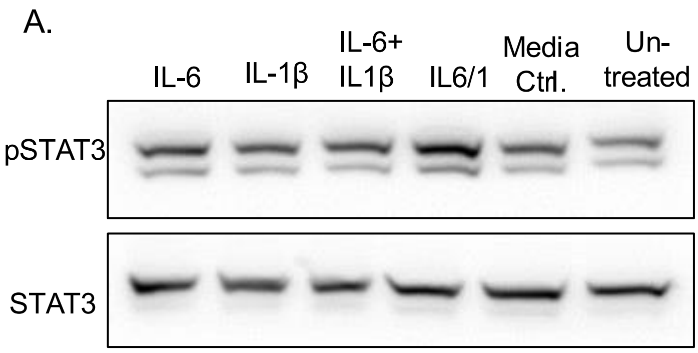


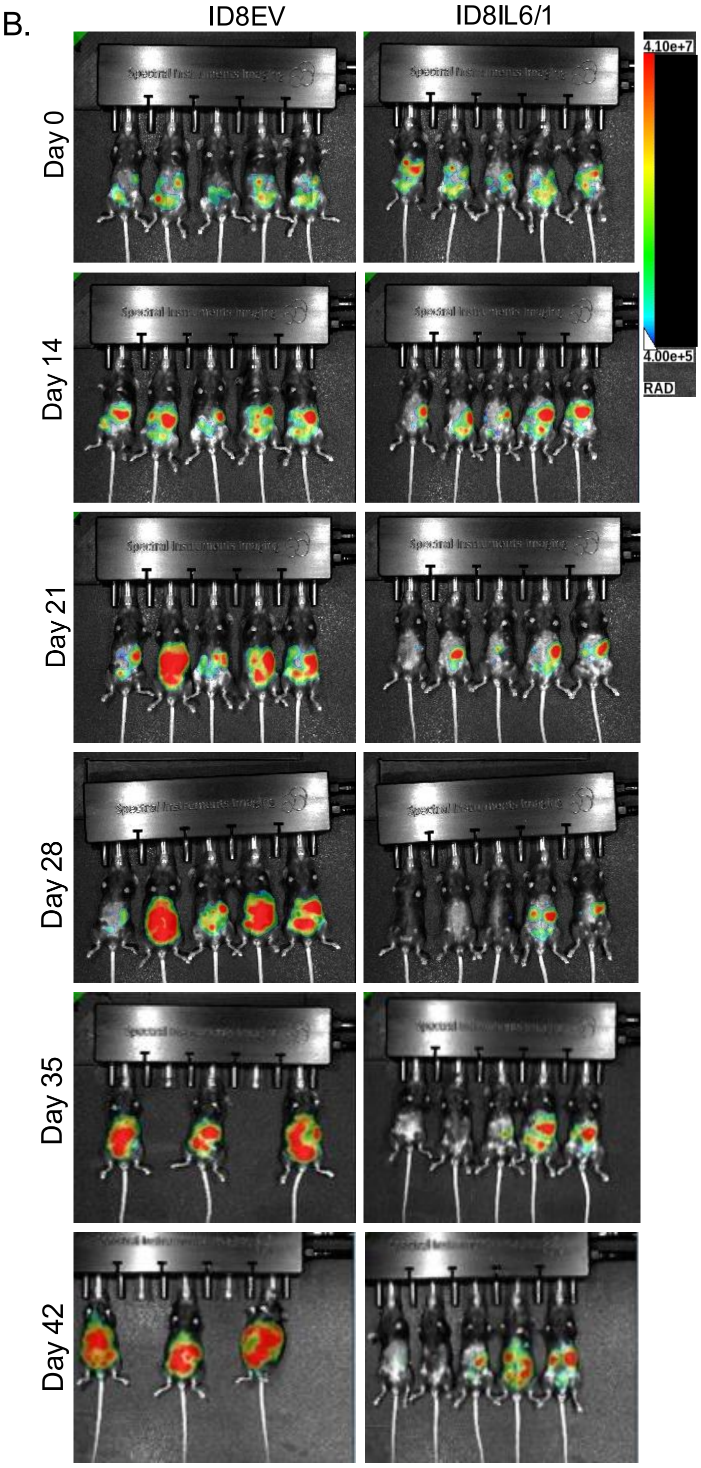


**Supplementary Figure 2:** (A) Human PBMCs derived CD3 T cells were induced to proliferate in the presence of equivalent amount of IL-6. IL1β, IL-6 + IL1β, IL6/1. After 72 h of culturing, cells were collected, lysed and subjected to western blot. (B) Tumor growth was quantitatively analyzed using bioluminescence (BLI) signals obtained from LAGO spectrum at a week after tumor inoculation and subsequent weeks. Longitudinal images of a representative mouse from each treatment group were presented from a week after tumor inoculation before treatment and subsequent weeks.
